# Functional binding of PD1 ligands predicts response to anti-PD1 treatment in cancer patients

**DOI:** 10.1101/2023.02.09.527671

**Authors:** Bar Kaufman, Orli Abramov, Anna Yevko, Daria Apple, Mark Shlapobersky, Yariv Greenshpan, Ruthy Shaco-Levy, Keren Roubinov, Alejandro Liboff, Moshe Elkabets, Angel Porgador

**Author notes:** Corresponding Authors: Angel Porgador Ph.D., The Shraga Segal Dept. of Microbiology, Immunology and Genetics, Ben-Gurion University of the Negev, Beer-Sheva, 84105, Israel. Moshe Elkabets Ph.D., The Shraga Segal Dept. of Microbiology, Immunology and Genetics, Ben-Gurion University of the Negev, Beer-Sheva, 84105, Israel.

## Abstract

Accurate predictive biomarkers of response to immune checkpoint inhibitors (ICIs) are required for better stratifying cancer patients to ICI treatments. Here, we present a new concept for a bioassay to predict the response to anti-PD1 therapies, which is based on measuring the binding functionality of PDL1 and PDL2 to their receptor, PD1. In detail, we developed a cell-based reporting system, called the Immuno-checkpoint Artificial Reporter with overexpression of PD1 (IcAR-PD1) and evaluated the PDL1 and PDL2 binding functionality in tumor cell lines, patient-derived xenografts, and in fixed-tissue tumor samples obtained from cancer patients. In a retrospective clinical study, we found that the functionality of PDL1 and PDL2 predicts response to anti-PD1, and functionality of PDL1 binding is a more effective predictor than PDL1 protein expression alone. Our findings suggest that assessing the functionality of ligand binding is superior to staining of protein expression for predicting response to ICIs.

**Teaser:** Positive clinical response of cancer patients to anti-PD1 therapy can be predicted by measuring the binding activity of PDL1 and PDL2.

## Introduction

Antibodies that have shown the most potent anti-tumor activity in cancer patients are those that block the immunomodulator, programmed cell death protein 1 (PD1) receptor (*1–4*), and its ligand, PDL1(*5, 6*) (i.e., pembrolizumab and nivolumab for PD1 and durvalumab for PDL1), but there is currently no single clinically validated biomarker that predicts the response to anti-PD1/PDL1 therapies in all cancer types (*7, 8*). As a result, different methods are used for predicting the response to antibody therapies for specific cancer types. For example, IHC for staining genomic instability of microsatellite instability (MSI) (*9*) gave predictive values of 39.6% for CRC and 34.3% for non-CRC cancers (*10*). Currently, the most widely used methodology for selecting cancer patients for anti-PD1/PD-L1 therapies is IHC staining for PDL1 (*11, 12*), which has an overall predictive value of 28.9% (*13*). It should be noted, however, that all current diagnostic assays and technologies, especially staining for PDL1 –being based on gene or protein abundances – cannot provide any insight into protein function and activity.

The function of immunomodulators is critical for cytotoxic immune-cell activity, as anti-tumor lytic activity is determined by the balance between immunomodulators that activate and suppress signals. Specifically, the suppression of the activity of cytotoxic T-cell lymphocytes is regulated by PD1 and its interaction with its mature ligands, PDL1 and PDL2 (PD1 ligands) (*14–16*). The maturation of these PD1 ligands requires post-translation modifications (*17–20*) and trans-localization into the cell membrane (*20–23*). Since anti-PD1 therapies block the interaction of PD1 with the mature forms of PD1 ligands, we posited that quantification the functionality of PD1 ligands binding to their receptor PD1 in tumor samples would be superior to PDL1 staining for predicting the response to anti-PD1 treatment.

Here, we elaborate on this novel concept that the functionality of PD1 ligands binding can be exploited to predict the response of cancer patients to anti-PD1 therapies. For this purpose, we first developed an engineered live-cell-based system, which we designated Immune-cell Artificial Reporter overexpressing PD1 (IcAR-PD1), that ‘reports’ the functionality of PD1 ligands binding. We then confirmed that IcAR-PD1 accurately quantifies PDL1 or PDL2 binding on the surface of tumor cells and validated its utility in fresh and formalin-fixed patient-derived xenografts. Thereafter, we developed a scoring system that calculates the functionality of each ligand (separately and together) in blocking PD1 and PDL1 with clinically approved anti-PD1 and anti-PDL1 therapies. Lastly, in a retrospective clinical trial we analyzed tumor samples from cancer patients treated with anti-PD1 therapies and demonstrated that IcAR-PD1 scoring is superior to PDL1 staining in predicting the response to anti-PD1 therapies. Overall, the results of this study indicate that our innovative approach, which quantifies the functionality of PDL1 and PDL2 binding to PD1, should be further developed as a predictive biomarker for the response to anti-PD1 therapy. The results also provide a strong rationale for studying the activity of other immunomodulatory ligands in predicting the response to other ICIs.

## Results

### Generation and characterization of the IcAR-PD1 cell line

To generate an immune-cell reporter line with the ability to quantify the functionality of PD1 ligands binding, we transduced BW-5147 murine thymoma cells (termed BW cells) to express recombinant human PD1 (rhPD1) fused to the murine CD3ζ chain (Fig. 1A and fig. S1). We verified that the modified BW cells expressed the chimeric protein by staining the cells with anti-hPD1 antibodies and analyzed the expression levels using flow cytometry (Fig. 1B). These ζ-fused PD1-expressing cells were termed IcAR-PD1. When IcAR-PD1 cells were introduced in culture wells pre-coated with the clinically approved anti-PD1 antibodies, pembrolizumab and nivolumab, the zeta chain was activated, resulting in the production and secretion of interleukin-2 (IL-2) (Fig. 1C and D). Notably, pre-coating the wells with an isotype control antibody (hIgG4) or with the clinically approved anti-PDL1 antibodies, durvalumab and avelumab, did not induce detectable IL-2 secretion by the IcAR-PD1 cells.

**Fig. 1.**
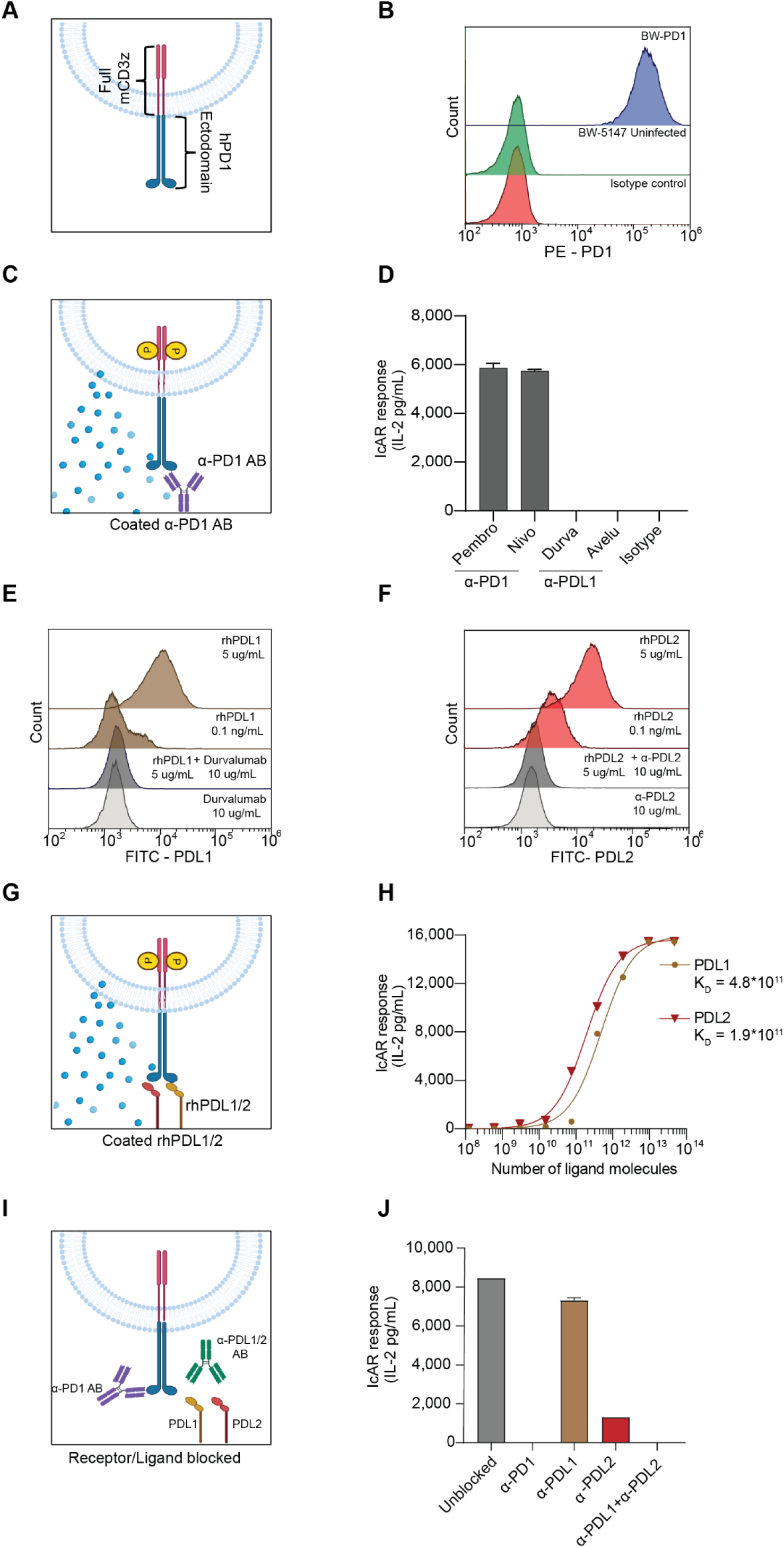
IcAR-PD1 expresses functional hPD1 receptors. (**A**) Schematic representation of the IcAR system, in which the ectodomain of the human PD1 receptor is fused to the full murine CD3ζ chain. Flow cytometry shows surface expression of hPD1 in the IcAR-PD1 cell line (top panel) vs. hPD1 expression in wild-type BW-5147 cells (middle panel) and isotype control (bottom panel). (**C**) Schematic representation of the functional activation of the IcAR-PD1 cell line in culture plates with wells precoated with anti-PD1 antibodies. Interaction between IcAR-PD1 cells and the anti-PD1 antibody resulted in the secretion of murine IL-2. (**D**) Stimulation of IcAR-PD1 cells with the anti-PD1 antibodies, pembrolizumab (Pembro) and nivolumab (Nivo), and anti-PDL1 antibodies, durvalumab (Durva) and avelumab (Avelu), or isotype control (human IgG4). (**E**) Flow cytometry analysis of IcAR-PD1 cells with recombinant Fc-hPDL1. (**F**) Flow cytometry analysis of IcAR-PD1 cells with recombinant Fc-hPDL2. (**G**) Schematic representation of the response of IcAR-PD1 cells to different ligands. (**H**) Response of IcAR-PD1 to different amounts of plastic-bound recombinant Fc-PDL1 and recombinant Fc-PDL2. (**I**) Schematic representation showing IcAR-PD1 cells blocked by different antibodies. (**J**) IcAR-PD1 cells were incubated with equimolar concentrations of rPDL1 and PDL2 without antibodies (unblocked) or with blocking antibodies: anti-PD1 antibody (pembrolizumab), anti-PDL1 antibody (durvalumab), anti-PDL2 antibody, and combination of anti-PDL1 and anti-PDL2 antibodies.

To test whether the hPD1 ectodomain expressed on the IcAR-PD1 cells could bind to PD1 ligands (*16*), we treated the cells with recombinant human PDL1-Fc and PDL2-Fc (rhPDL1 and rhPDL2, respectively) and then stained them against the Fc domain of the recombinant ligands for flow cytometry analysis (Fig. 2E and F). To confirm the specific binding of rhPDL1 and rhPDL2 to PD1 on the IcAR-PD1 cells, we neutralized ligand binding with the relevant antibodies and observed a reduction in ligand binding (Fig. 2E and F). We found that staining with lower concentrations of the recombinant ligands resulted in better staining by rhPDL2 than by rhPDL1, which is in line with the higher affinity of PDL2 for PD1, as previously reported (29).

**Fig. 2.**
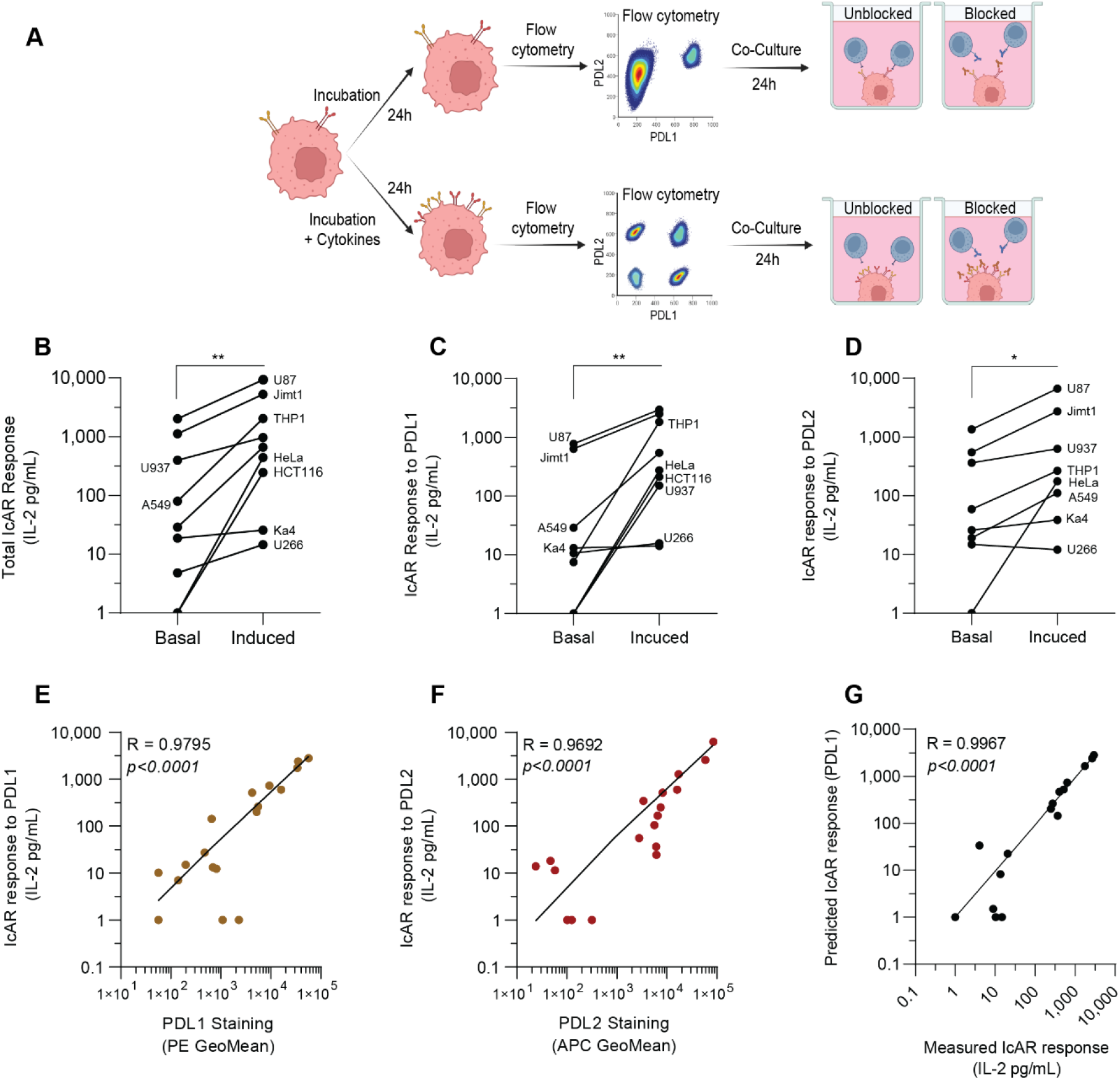
IcAR-PD1 can discriminate between PDL1 and PDL2 in cancer cell lines. (**A**) Expression levels of PDL1 and PDL2 were measured for each cell line (basal and cytokine-induced) using flow cytometry; the IcAR functional assay was performed on both unblocked and blocked ligands using anti-PDL1, anti-PDL2 and anti-PD1 antibodies. (**B**) The IcAR response to nine different cell lines increased upon cytokine induction (paired t-test, *P =*0.002). (**C**) The IcAR response to PDL1 in response to nine different cell lines increased upon cytokine induction (paired t-test, *P =*0.0039) (**D**) The IcAR response to PDL1 in response to nine different cell lines increased upon cytokine induction (paired t-test, *P =*0.0156). (**E**) Correlation between the IcAR-PDL1 response and PDL1 surface expression on tumor cells (n=18). (**F**) Correlation between the IcAR-PDL2 response and PDL3 surface expression on tumor cells (n = 18). (**G**) Prediction of response to PDL1 calculated by subtracting the IcAR response post PDL1 blockade from the overall IcAR response, without the use of an anti-PDL2 antibody (n=18). Differences were considered to be statistically significant at a two‐sided *P < 0.05, **P < 0.01. PE - phycoerythrin, APC - allophycocyanin, GeoMean - geometric mean fluorescence intensity.

We then investigated the ability of rhPDL1 and rhPDL2 to activate IcAR-PD1 cells to secrete IL-2 by exposing the cells to increasing concentrations of rhPDL1 or rhPDL2 (Fig. 1H). The calculated binding affinity of each ligand, based on the secretion of IL-2 in response to varying ligand concentrations, indicated that PDL2 has a higher affinity (2.5-fold) for PD1 (Kd 4.82×1011) than PDL1 (Kd 1.94×1011). Lastly, we demonstrated that anti-PD1 or anti-ligand antibodies blocked IL-2 secretion from IcAR-PD1 cells exposed to well-bound rhPDL1 and rhPDL2 (ligand ratio of 1:1) (Fig. 1J). The anti-PD1 antibody, pembrolizumab, completely blocked IL-2 secretion, while the anti-PDL2 antibody was more effective than the anti-PDL1 antibody (durvalumab) in blocking IL-2 secretion. These results indicate that IcAR-PD1 could accurately identify and distinguish between the binding of PDL1 and PDL2 to PD1 and could accurately measure the functional binding of each ligand to understand the contribution of each to PD1 activation.

### IcAR-PD1 response to cancer cells is dependent upon both PDL1 and PDL2

To explore the ability of IcAR-PD1 to quantify the functionality of PD1 ligands binding in tumor cells, we first needed to establish the optimal co-culture conditions for inducing IL-2 secretion. We used A549 cells treated with IFNγ plus TNFα as target cells and co-cultured them with IcAR-PD1 in 96-well plates (fig. S2). We found that the minimal number of IcAR-PD1 cells required to detect as few as 700 target cancer cells is 10^5^. We then investigated whether IcAR-PD1 could recognize, discriminate, and quantify the binding functionality of PDL1 and PDL2 expressed on the surface of nine cancer cell lines under normal conditions and following stimulation with IFNγ and TNFα (shown schematically in Fig. 2A). We used a co-culture of tumor and IcAR-PD1 cells in which PDL1 and PDL2 were blocked and assessed the functionality of their binding in each tumor cell line. Using flow cytometry, we measured the surface expression of PD1 ligands and found that their abundance increased after cytokine stimulation in all nine cell lines (fig. S3 and table S1 present the complete information for each cell line). The IcAR-PD1 response was also enhanced in co-culture tumor cells stimulated with IFNγ and TNFα compared to non-stimulated cells (Fig. 2B). This profiling of the nine cell lines showed that exposure to IFNγ plus TNFα induced upregulation and activation of PDL1 and PDL2. We used antibodies to differentiate between the two PD1 ligands and observed their functional binding in nine cell lines. We found that functional binding of PDL1 (*P =*0.0039) and PDL2 (*P =*0.0156) was significantly enhanced in most cell lines (Fig. 2C and 2D). By integrating the information for the nine cell lines for the expression levels of cell-surface PD1 ligands with the results for PD1 ligands activity upon cytokine stimulation, we were able to calculate the correlation between cell-surface expression of PDL1 or PDL2 and the IcAR-PD1 response. We found strong correlation (R = 0.951, P < 0.0001) between the cell surface expression of PDL1 and the IcAR-response to PDL1 in the presence of an anti-PDL2 antibody (Fig. 2E). Similarly, we observed strong correlation (R = 0.969, P < 0.0001) between PDL2 cell-surface staining and the IcAR response to PDL2 (Fig. 2F). We then examined whether we could calculate the functional binding of PDL1 and PDL2 in tumor cells by neutralizing only PDL1. To do so, we subtracted the IcAR-PD1 response levels post-treatment with anti-PDL1 antibody (response to PDL2) from the total IcAR response (unblocked) to obtain a “predicted IcAR response to PDL1”. We found a strong association (R = 0.997, P < 0.0001) between predicted and measured responses to PDL1 in the nine different cell lines (Fig. 2G), indicating that the activation capacity of both ligands can be predicted by measuring the response to only one ligand. These results demonstrate the potential of IcAR-PD1 as a tool for predicting the functionality of PD1 ligands binding in a live-cell system.

### IcAR-PD1 can measure PDL1 and PDL2 in fresh and fixed tumor tissues

To evaluate the IcAR-PD1 response in fresh tumors, we used five patient-derived xenografts (PDXs) obtained from patients with lung or head and neck cancer (table S2). We used a single-cell suspension of each PDX to determine the expression of the ligands by flow cytometry (Fig. 3B) and to determine the functional binding of PD1 ligands upon co-culture with IcAR-PD1 (Fig. 3C); the scheme for testing fresh PDX samples is shown in Fig. 3A. PDX numbers 1, 2, and 3 showed near-absolute PDL1 expression and activation levels, while PDX numbers 4 and 5 expressed both PDL1 and PDL2. In PDX number 4, the PDL2-based IcAR response was near absolute for PDL2, as only 9% of the IcAR response was attributed to PDL1. In contrast, PDX number 5 showed a mixed IcAR response to PDL1 and PDL2, similar to the surface expression pattern of PD1 ligands.

**Fig. 3.**
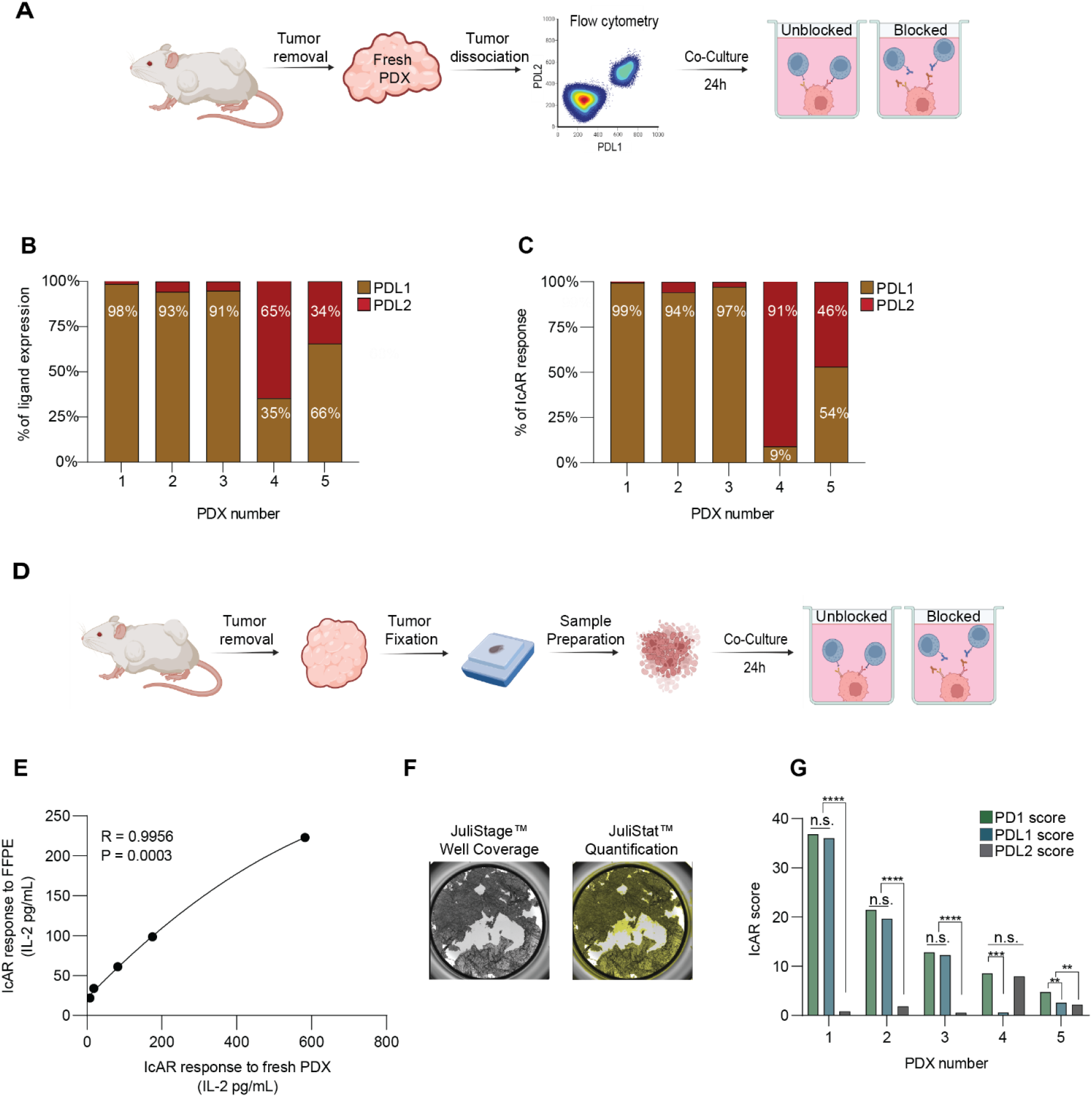
IcAR response to fresh and fixed PDXs and generation of an IcAR score. (**A**) Schematic representation of the IcAR assay protocol. Following flow cytometry determination of surface-expression levels of PDL1 and PDL2 and co-expression of PDL1 and PDL2 (double) in fresh PDXs (extracted from mice), the IcAR functional assay was performed on both unblocked ligands and ligands blocked using anti-PDL1, anti-PDL2 and anti-PD1 antibodies. (**B**) PDL1 and PDL2 levels were determined in five PDXs, and the assay results are presented as percent of surface expression of each ligand and of co-expression (double). (**C**) IcAR functional assay was performed for each PDX using anti-PD1, anti-PDL1 and anti-PDL2 antibodies; results are displayed as percent of response to each ligand out of the total IcAR response (unblocked). (**D**) IcAR functional assay was performed for each FFPE tissue sample using anti-PD1, anti-PDL1 and anti-PDL2 antibodies; results are displayed as percent of response to each ligand out of the total IcAR response (unblocked). (**E**) Correlation between IcAR response (namely, IL-2 levels) from fresh and fixed PDXs (n = 5). (**F**) Example of surface coverage of a well, measured with a JuliStage™ cell recorder, to produce normalized response per surface area. (**G**) Different PDXs give different scores, depending on the functionality of binding of each ligand. (\*\**P <* 0.01; \*\*\*\**P <* 0.0001).

Since obtaining fresh biopsies from cancer patients is logistically challenging, we sought to develop the IcAR-PD1 technology to measure PDL1 and PDL2 activity in formalin-fixed paraffin-embedded (FFPE) tissue samples. We optimized an antigen retrieval protocol for 5-micron slices to establish a consistent response of IcAR-PD1 (Fig. 3D). We then used the five PDXs to compare the IcAR-PD1 response in fresh and FFPE tumors (Fig. 5E). We found that even though the overall IcAR response levels were lower for fixed tissues, there was strong correlation between the results for fresh and fixed PDXs (Pearson coefficient 0.995; *P =*0.0003), indicating the potential of IcAR-PD1 to measure PDL1 and PDL2 activity in FFPE samples.

### Development of an IcAR score for evaluating PD1 ligands activity in FFPE samples

To accurately assess the functionality of PD1 ligands binding in FFPE samples, we generated a Formula (1) that gives a score that incorporates: (i) the response of IcAR-PD1 (in terms of IL-2 levels) with and without blocking anti-PD1 or anti-PDL1 antibodies, (ii) the amount of tissue (quantification of the coverage area of cell aggregates in each well), and (iii) normalization to a maximum response induced as a positive control (to normalize data between experiments).

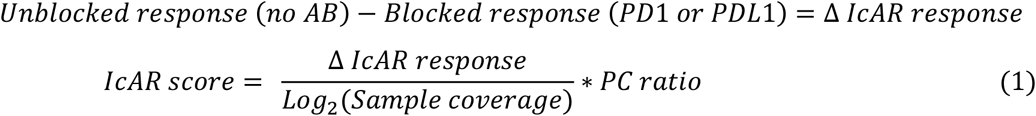

where IL-2 measurements of each sample well were divided by log2 of sample coverage, as measured and analyzed by JuLiStat™ to produce IL-2 signal per log_2_(mm^2^). Thereafter, the IcAR score for each sample well was calculated for each antibody separately, using the above formula.

The results obtained from our formula generate an IcAR score instead of an IcAR response. Since the IcAR response is assessed by blocking the signals with anti-PD1 or anti-PDL1 antibodies, we can differentiate between PDL1 and PDL2 responses and generate a total IcAR-PD1 score (termed the PD1-score) or a PDL1-based IcAR-PD1 score (termed the PDL1-score). Since the activation capacity of the two ligands can be calculated using only anti-PDL1 antibodies (Fig. 2G), we also generated a scoring formula for PDL2, which we called the PDL2-score. Fig. 3G shows the PD1, PDL1, and PDL2 scores for all five fixed PDX samples. The PD1 and PDL1 scores were similar for PDX numbers 1-3 and reflected ∼100% expression of cell-surface PDL1. There was a significant difference between PD1 and PDL1 scores in PDX numbers 4 and 5, as the signal for PD1 activation in IcAR-PD1 cells is also induced by PDL2.

### Description of the patient cohort for which PD1 and PDL1 scores were generated

To investigate whether IcAR-PD1 scores can predict the clinical response to anti-PD1, we collected FFPE tumor samples from 42 patients who had been treated with anti-PD1 therapy. The distribution of the samples by cancer type is shown in Fig. 4A. Most of the samples were derived from NSCLC patients (20 samples, 57%), the majority with adenocarcinoma (13) and the remainder (7) with squamous cell carcinoma. For the remaining 22 samples, transitional cell carcinoma accounted for 9, renal cell carcinoma for 9, melanoma for 2, and head and neck squamous cell carcinoma for 2. Of the entire cohort, 23 patients were classified as non-responders, and their disease was classified as progressive or stable, while 19 of the patients were classified as responders, and their clinical response was classified as a partial response or a complete response (Fig. 4B). There were no significant differences between the responders and non-responders in terms of age (Fig. 4C), gender, white blood cell count, or lymphocyte count (table 1), but for overall survival there was a significant difference between responders and non-responders (Fig. 4D; log-rank Kaplan-Meier plot, *P =*0.0002).

**Table 1.** Clinical Cohort. The table presents statistics regarding the differences between responders and non-responders. Statistical significance was determined by a t-test for parametric values, or Chi-square for non-parametric values. * Percentage of total population Abbreviations: RCC = renal cell carcinoma, TCC = transitional cell carcinoma, H&N = head and neck cancer. Chemo = chemotherapy, Pembro = pembrolizumab (anti-PD1), Ipi = ipilimumab (anti-CTLA4), Nivo = nivolumab (anti-PD1), Anti-PD1 = nivolumab/pembrolizumab. WBC = white blood cells.

|  |  | **Total** | **Responders** | **Non-responders** | **P** |
| --- | --- | --- | --- | --- | --- |
| **N** |  | 42 | 19 (45.23%)* | 23 (54.77%)* |  |
| **Average Age - Years** |  | 69.56<br>95%CI (66.51 – 72.59) | 67.42<br>95%CI (62.74 – 72.10) | 71<br>95%CI (67.13 – 75.48) | 0.8619 |
| **Sex** | Female | 16 (38.1%)* | 7 (36.84%) | 9 (39.13%) | 0.8792 |
|  | Male | 36 (61.9%)* | 12 (63.16%) | 14 (60.77%) |  |
| **Stage** | 3 | 2 (4.76%)* | 1 (5.26%) | 1 (4.35%) | 0.8897 |
|  | 4 | 40 (95.24%)* | 18 (94.74%) | 22 (95.65%) |  |
| **Pathology** | RCC | 9 (21.43%)* | 2 (10.53%) | 7 (30.43%) | 0.3138 |
|  | TCC | 9 (21.43%)* | 5 (26.32%) | 4 (17.39%) |  |
|  | Lung | 20 (47.62%)* | 9 (47.37%) | 11 (47.83%) |  |
|  | H&N | 2 (4.76%)* | 2 (10.53%) | 0 |  |
|  | Skin | 2 (4.76%)* | 1 (5.26%) | 1 (4.34%) |  |
| **ICI treatment** | Anti-PD1 | 7 (16.67%)* | 4 (21.05%) | 3 (13.04%) | 0.2843 |
|  | Anti-PD1 + Chemo | 28 (66.67%)* | 14 (73.68%) | 14 (60.88%) |  |
|  | Pembro + axitinib | 3 (7.14%)* | 0 | 3 (13.04%) |  |
|  | Ipi + Nivo | 4 (9.52%)* | 1 (5.26%) | 3 (13.04%) |  |
| **Pretreatment blood count** | WBC (cells/dL) | 8211<br>95%CI (7386 – 9037) | 8773<br>95%CI (7537 – 10001) | 7280<br>95%CI (6666 – 8976) | 0.5292 |
|  | Lymphocytes (cells/dL) | 1992<br>95%CI (1633 – 2352) | 2220<br>95%CI (1563 – 2843) | 1846<br>95%CI (1396 – 2296) | 0.5337 |
| **Mortality** |  | 23 (54.76%) | 7 (36.84%) | 15 (65.22%) | 0.0669 |
| **Average overall survival (days)** |  | 679<br>95%CI (535 – 823) | 823<br>95%CI (606 – 1041) | 555<br>95%CI (363 – 746) | 0.0546 |
| **IcAR score** | PD1 | 7.24<br>95%CI (5 – 9.5) | 13.49<br>95%CI (10.45 – 16.54) | 2.07<br>95%CI (1.422 – 2.736) | <0.0001 |
|  | PDL1 | 6.35<br>95%CI (4.228 – 8.848) | 11.95<br>95%CI (8.797 – 15.11) | 1.73<br>95%CI (1.075 – 2.385) | <0.0001 |

**Fig. 4.**
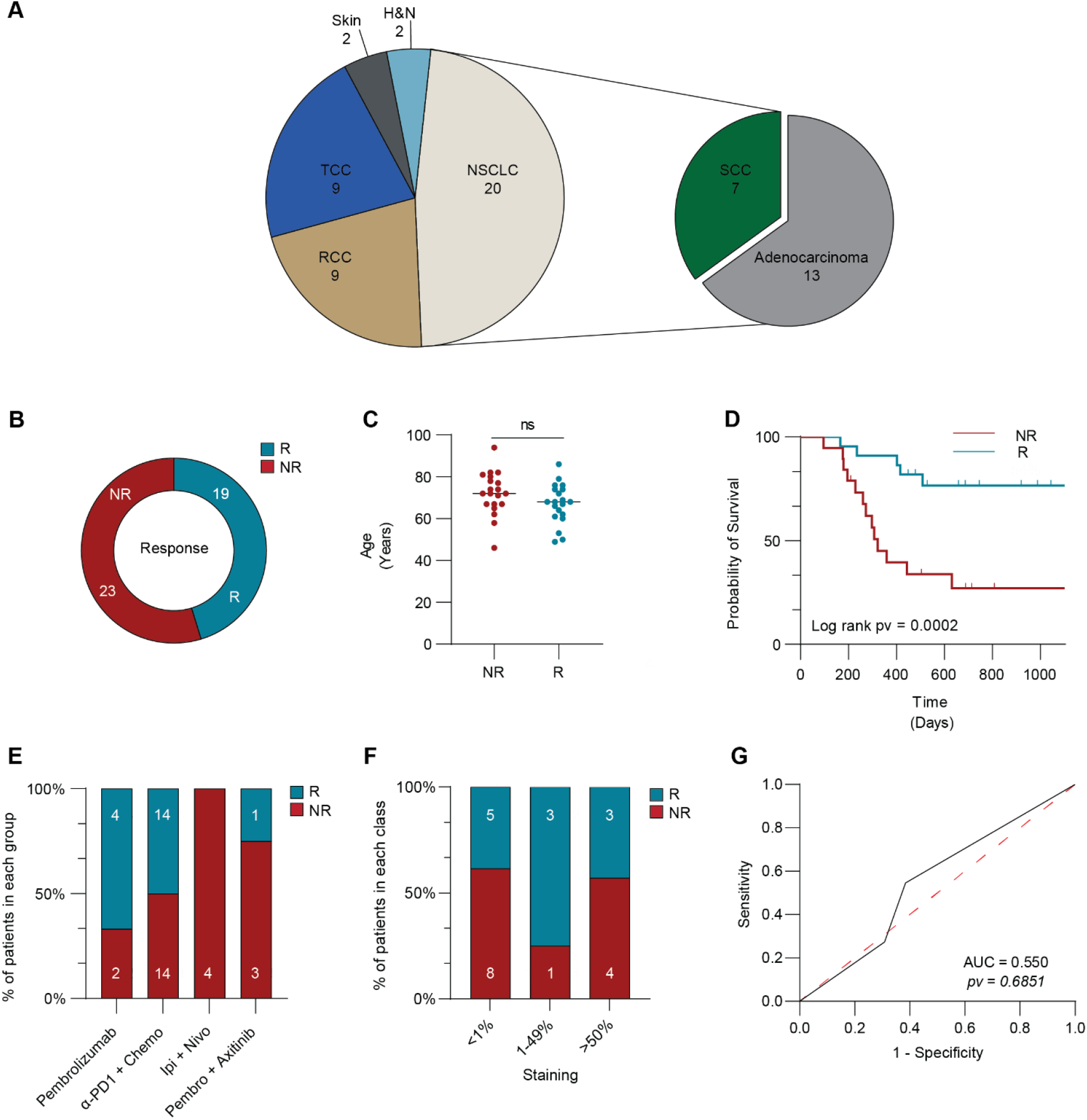
Description of the patient cohort. (**A**) Pie chart showing the distribution of patients with different cancer types (n = 42). H&N - head and neck cancer, NSCLC - non-small cell lung carcinoma, RCC-renal cell carcinoma, TCC - transitional cell carcinoma, SCC - squamous cell carcinoma. (**B**) Distribution of cohort into two categories, responders (R) and non-responders (NR). (**C**) The difference in age distribution between responders (R; blue) and non-responders (NR; red) in the cohort (n = 42) is not significant (*P =*0.121). (**D**) Survival curves showing significant differences in survival time between responders and non-responders (log-rank *P =*0.0063, n = 42). (**E**) Distribution of treatment groups between responders and non-responders (n = 42). Ipi - ipilimumab, Nivo - nivolumab, Pembro – pembrolizumab. (**F**) Distribution of PDL1 status between responders and non-responders (n = 24). (**G**) Logistic regression of the predictive value of PDL1 for a patient’s clinical response is shown as a ROC curve: Red line depicts a random model, black line depicts PDL1 staining (AUC = 0.53, *P =*0.79, n = 29).

**Fig. 5.**
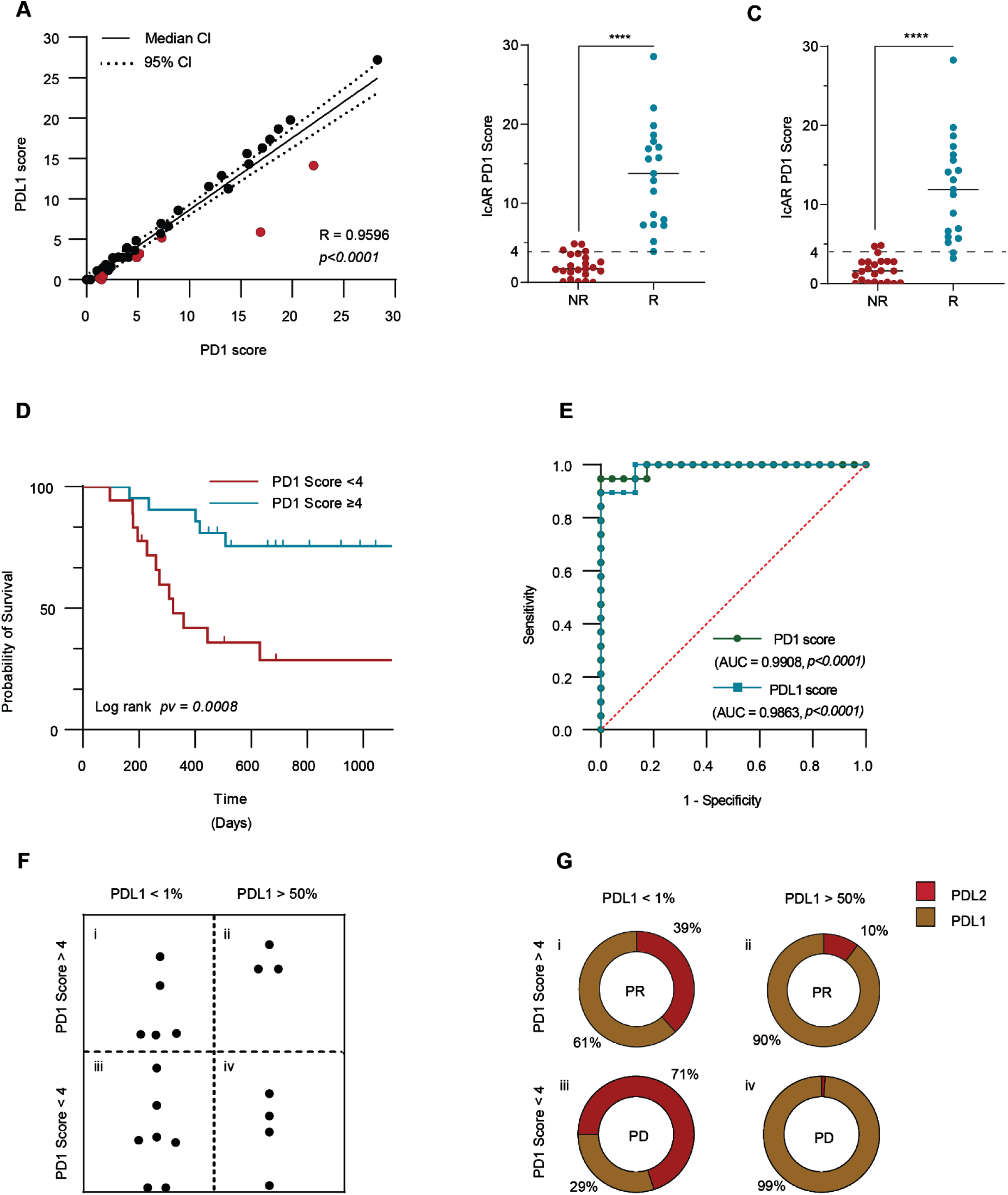
PD1 and PDL1 score can distinguish between responders and non-responders. (**A**) Correlation between PD1 score (X) and PDL1 score (Y) for each patient (Pearson R = 0.9596, *P <* 0.0001, n = 42); patients with significantly low PDL1 scores are represented by red dots. (**B**) IcAR-PD1 score in responder (R) and non-responder (NR) groups (\*\*\*\**P* < 0.0001, n = 42). (**C**) IcAR-PDL1 scores in responder (R) and non-responder (NR) groups (\*\*\*\**P <* 0.0001, n = 42). (**D**) Survival curve of patients with IcAR score ≥ 4 and patients with an IcAR score < 4 (log-rank *P =*0.0008, n = 42). (**E**) Logistic regression of the predictive value of PD1 and PDL1 scores for a patient’s clinical response is shown as a ROC curve: red line depicts a random model, green line depicts PD1 score (AUC = 0.990, *P <* 0.0001, n = 42), blue line depicts PDL1 score (AUC = 0.9863, *P <* 0.0001). (**F**) Comparison between PD1 score (≥ 4 and < 4) and PDL1 staining. (**G**) Ligand distribution and prognosis in four patients, one from each of the groups shown in (F). The outer circle shows IcAR response patterns of four patients, and inner circle shows the patient’s clinical response, where PR = partial response and PD = progressive disease.

The proportion of responders and non-responders in the different treatment categories is shown in Fig. 4E. The most common treatment was a combination of anti-PD1 with chemotherapy (28/42 patients; 66.7% of the cohort). PDL1 staining and pathological evaluation were performed for 24 of the 42 samples. Pathological evaluation of PDL1 staining was divided into three categories: negative (< 1%); positive (1-49%); and strongly positive (≥ 50%). Among the 24 tumors for which we had a PDL1 staining evaluation, 13 were PDL1 negative; for these tumors, 5/13 (38.5%) were classified as anti-PD1 sensitive (patients responded to therapy), and 8 were classified resistant (patients did not respond to treatment) (Fig. 4F). Of the seven patients classified with PDL1 strongly positive tumors, 3 (42.8%) were classified as anti-PD1 sensitive and 4 (57.2%) as resistant. To examine the link between the pathological evaluation of PDL1 staining and the clinical response of each of the 29 patients, we performed a simple logistical regression. The results, shown in Fig. 4G as an area under the ROC curve (AUC), indicate a non-significant association (*P =*0.747) (AUC = 0.535), with a negative predictive value of 45.5% and positive predictive value of 47.06%.

### PD1 and PDL1 scores correlate with the clinical response to anti-PD1 treatment

To determine the functionality of PD1 ligands in cancer patients, we co-cultured IcAR-PD1 cells with 5-micron slices of tumors from 42 cancer patients who had undergone anti-PD1 treatment. We first prepared the samples by deparaffinizing them and performing antigen retrieval. We then seeded the samples in a 96-well plate and determined the well coverage for each sample. We added IcAR-PD1 cells to each well and cultured them for 18 hours in the presence of anti-PD1 or anti-PDL1 antibodies. After the culture period, we measured the levels of mIL-2 in the culture medium using ELISA. We calculated both the PD1 and PDL1 scores for each patient (Fig. 5A). In general, most patients had similar PD1 and PDL1 scores, but some patients had significantly lower PDL1 scores, suggesting a higher level of PDL2 involvement.

When stratifying the IcAR-PD1 scores by response to treatment, we found that the PD1 and PDL1 scores exhibited similar patterns (Fig. 5B and 5C), with lower scores being less likely to respond to anti-PD1 treatment. For both the PDL1 and PD1 scores, there was a significant difference in values between responders and non-responders (P<0.0001). The difference in the PD1 score between responders and non-responders was reflected in the patient overall survival, as shown by a Kaplan-Meier plot in Fig. 5D; for patients with a PD1 score of 4 or higher (defined by 95% confidence interval of non-responder: mean = 2.07 ± 2SD), the overall survival was significantly higher than that for patients with a PD1 score of less than 4 (log-rank *P =*0.0008). The mean OS of responses was 823 days (95%Cl 606–1041 days), while for the non-responders, the mean OS was 555 days (95%Cl 363–746 days). We also calculated the predictive value of PD1 and PDL1 scores using multiple logistic regression, with AUCs of 0.9908 and 0.9863, respectively (Fig. 5E). The positive and negative predictive values for the PD1 score were 100% and 95.8%, respectively, while the positive and negative predictive values for the PDL1 score were 91.3% and 89.4%, respectively.

When we compared the PD1 score to the pathological grading of PDL1 in tumors as negative (<1%) or strongly positive (>50%) (Fig. 5F), we found four different groups of tumors: (i) patients with negative PDL1 staining and high PD1 scores, (ii) patients with strong PDL1 staining and high PD1 scores, (iii) patients with negative PDL1 staining and low PD1 scores, and (iv) patients with strong PDL1 staining and low PD1 scores. To further explore the differences between the samples, we analyzed the percent of IcAR response to PDL1 and PDL2 within each staining group (Fig. 5G and fig. S4). For example, for a patient belonging to group iv (strong PDL1 staining and low PD1-IcAR score), the ligand was predominantly PDL1 (99%), and the patient did not respond to treatment. In contrast, for a patient belonging to group i (negative PDL1 staining but high PD1-IcAR score), the ligand distribution was also PDL1 dominant, but with 61% PDL1 and 39% PDL2, and the patient exhibited a partial response to therapy.

## Discussion

Immune checkpoint inhibitors (ICIs) have been shown to extend the survival of some cancer patients in medical oncology (*24*). However, several challenges exist with using ICIs to treat cancer, including low response rates (*25, 26*), potential for severe adverse events(*27, 28*), and high costs (*29*). Identifying patients who are more likely to benefit from ICI treatment could save lives and improve patient outcomes(*30*). Current predictive methodologies, such as PDL1 staining, microsatellite instability (MSI) analysis, and tumor mutation burden (TMB) analysis, are insufficient and other technologies and assays are needed to improve their predictive value (*12, 31, 32*). For example, IHC for staining of MSI (*9*) predictive value is 34.3% for non-colorectal cancers (*10*). Genomic sequencing of the tumor sample and quantification of TMB(*33*) can be used for predicting response in non-small cell lung cancer, colorectal carcinoma, and transitional cell carcinoma (*34, 35*). Machine learning and artificial intelligence analyses of multi-omics data have also shown promising predictive values for non-small cell lung cancer (*36–38*). However, methods based on immunohistochemistry staining for ligand abundancy remains the most commonly used at present (*11*).

Specifically for PD1/PDL1 immunotherapy, the implementation of PDL1 staining as a predictive marker is not satisfactory, with an overall predictive value as low as 28.9% (*12, 31, 32*). The reasons for this relatively low predictive value could include: (i) the challenge of generating appropriate accurate diagnostic monoclonal antibodies either to the non-proteinic PDL1 (*39, 40*) or to the mature glycosylated PDL1 that differentially affects the binding to the PD1 receptor (*41, 42*); (ii) the FDA-approved staining evaluation protocol (*43, 44*) divides PDL1 staining to 3 wide-ranging categories (i.e. positive PDL1 staining values range from 1% to 49% staining); (iii) the high inter-pathologist variability in the interpretation of staining results as an outcome of the wide range within the staining groups (*45*); and (iv) a lack of a FDA-approved staining methodology for evaluating the presence of PDL2, which is known to induce immune suppression via activation of PD1 (*46–48*). Here, we propose a new diagnostic technology that determines the binding functionality of PDL1 and PDL2, thereby overcoming, to some extent, the above challenges for three main reasons: (i) The assay is founded on cell-based reporting of a PDL1 and PDL2 binding-function and hence addresses the above-mentioned concerns regarding detection of multiple ligands, the effect of the PDL1 and PDL2 glycosylation state, and the influence of non-proteinic epitopes on IHC staining results. (ii) The co-culture test of IcAR-PD1 with the cancer tissue provides a continuous wide-range scale associated with PDL1 and PDL2 function, with higher sensitivity and specificity of ligand activity than IHC staining against PDL1. (iii) The functional test evaluates the ability of clinically approved anti-PD1/PDL1 therapies to prevent ligand activity and provides a scoring of the contribution of each ligand (PDL1 and PDL2 scores) in mediating immune escape for each individual patient.

Functional assays are used primarily for fresh tumor-tissue samples (*49*), but the logistics of sample collection, storage and shipping, including the high costs, complicate a large-scale routine diagnostic methodology. To overcome such obstacles and facilitate large-scale routine clinical trial diagnostics, we optimized the IcAR-PD1 to recognize PD1 ligands binding functionality in material derived from FFPE blocks. This optimization enabled us to perform a retrospective study to analyze 42 tumor samples of cancer patients treated with anti-PD1 therapy. A significant advantage of the assay is that it can be tailored by using the drug administered in a specific treatment to generate a score for that treatment. Specifically, we used FDA-approved anti-PD1 antibodies to quantify the total binding of PD1 ligands and found that IcAR-PD1-score was highly correlated with the clinical outcome of the anti-PD1 therapy. In addition to the total score for PD1 ligands, we could also calculate a score for each ligand separately, with the PDL1-score showing a strong correlation with the clinical benefit to patients treated with anti-PD1. It is important to note that for a few cancer patients, functional binding of PDL2 was higher than for PDL1, yet PD1-score remained highly indicative of their response to anti-PD1 therapy.

The relative contribution of PDL2 to the overall ligand abundance of each patient was uncovered when we compared the PD1-score with the pathological grading of PDL1. Our analysis showed for the first time that cancer patients with negative PDL1 staining can have functional PDL2 that activate IcAR-PD1, and such patients benefit from anti-PD1 treatment. Furthermore, we also showed that cancer patients with high PDL1 staining can have a none/low functional PDL1 and thus would not benefit from anti-PD1 treatment.

While today anti-PD1/PDL1 therapies are the major FDA-approved ICIs in medical oncology, it is very likely that in the coming years multiple checkpoint inhibitors will be approved by the FDA for many different targets (*50*). Some of these checkpoint receptors, such as TIM3 (*51*) and LAG3 (*52*), can bind to multiple ligands, and therefore pose a challenge for current predictive methods (*53*). It is likely that TMB and MSI staining, which are indicative of general tumor characteristics and can therefore indicate sensitivity to ICIs in general (*54*), will not be sufficiently informative to distinguish between sensitivity to different ICIs.

To summarize, our results indicate that quantification of the activity of immunomodulators can be used to predict the clinical response to immunotherapy, and thus the IcAR technology could be a valuable component of the toolbox of oncologists, supporting and prioritizing treatment regimens designed for precision cancer immunotherapy. Moreover, in a few years, there will probably be many FDA-approved ICIs that block LAG3, TIGIT, TIM3, and NKG2A (*50*), and decisions regarding specific ICI treatments will become even more challenging. Developing the IcAR methodology for other immunomodulators could potentially be used for stratifying cancer patients for treatment with ICIs.

## Materials and Methods

### Cell lines

Mouse BW5147 thymoma cells and transfectants were maintained in RPMI 1640 medium containing 10% (v/v) fetal calf serum (FCS), penicillin, streptomycin, glutamine, and sodium pyruvate (1 mM each). BW-derived reporter cell lines were maintained in RPMI medium containing selection antibiotics.

The A549, JIMT1, U-87MG, and HeLa cell lines were maintained in DMEM medium containing 10% (v/v) FCS, penicillin, streptomycin, glutamine, and sodium pyruvate (1 mM each). The U-937, THP-1, U-266, and Ka4/721.221 cell lines were maintained in RPMI-1640 medium containing 10% (v/v) FCS, penicillin, streptomycin, glutamine, and sodium pyruvate (1 mM each). The HCT-116 cell line was maintained in McCoy’s 5a medium containing 10% (v/v) FCS, penicillin, streptomycin, glutamine, and sodium pyruvate (1 mM each). All media and supplements were purchased from Biological Industries (Kibbutz Beit-Haemek, Israel).

### Production and selection of the functional IcAR-PD1 cell line

Cloned sequences encoding the human extracellular regions of several immune-checkpoint receptors fused to the murine CD3ζ chain were supplied by HyLabs (Israel). The extracellular portion of PD1 cDNA (NM_005018.2; (24F – 170V) was fused to mouse CD3ζ chain cDNA (NM_001113391.2; (31L-164R)) to produce PD1-ζ. The immune-checkpoint receptor sequence was cloned into pHAGE240. HEK-293T cells were transiently transfected with an hPD1-ζ-containing plasmid and lentiviral packaging plasmids, and retrovirus-containing supernatants were harvested, aliquoted and used for transduction of BW5147 thymoma cells. Following selection with puromycin (InvivoGen, CA, USA)/G418 (Sigma-Aldrich, MO, USA), stable transfectants were screened by flow cytometry using mouse anti human PD1 (cat: 621610, BioLegend, CA, USA) and isotype control using allophycocyanin (APC)-conjugated mouse IgG1(cat: 400120, BioLegend). BW5147 cells expressing a receptor fused to murine CD3ζ secrete IL-2 following activation of the receptor. To examine the effector:target ratio suitable for this assay, BW-PD1 cells were incubated with A549 cells induced with IFNγ and TNFα to express different immune-checkpoint ligands, followed by testing with a murine IL-2 commercial sandwich ELISA kit (BioLegend).

### Flow cytometry

For cell suspension preparation, cells were washed in RPMI medium, then with 1X PBS, re-counted, and plated at a concentration of 100,000 cells per well in a 96-well plate. Target cell lines were stained using phycoerythrin (PE)-conjugated mouse anti-human PDL1 antibody (cat: 329706, BioLegend) and APC-conjugated mouse anti-human PDL2 antibody (cat: 329608, BioLegend). PE-conjugated mouse IgG2b (cat: 402203, BioLegend) and APC-conjugated mouse IgG2a (cat: 981906, BioLegend) were used as isotype controls. The IcAR-PD1 cell line was stained using PE-conjugated mouse anti-human PD1 antibody (cat: 379210, BioLegend). PE-conjugated mouse IgG2b (cat: 402203, BioLegend) was used as the isotype control for the IcAR-PD1 cell line staining. Antibodies were added to a final concentration of 2 mg/mL and samples were incubated for 1.5 h on ice. Later, cells were washed twice with 1X PBS and resuspended with dead cell marker labeled by adding DAPI. All the samples were analyzed using Beckman CytoFLEX flow cytometer, which can detect up to 13 different fluorochrome-conjugated antibodies simultaneously. As a gating strategy, isotype-matched controls for each sample were analyzed to set the appropriate gates; representative data are reported in relevant figures. For each marker, samples were analyzed in triplicate. To minimize false-positive events, the number of double or single positive events detected with the isotype controls was subtracted from the number of double or single positive cells stained with corresponding antibodies (not isotype control), respectively. Cells expressing a specific marker were reported as a percentage of the total number of gated events (double or single positive).

### Cell co-culture

All cell lines were seeded in flat 96-well plates at a density of 0.25 × 10^5^ cells/well and cultured for 24 h in the absence or presence of human recombinant IFNγ and TNFα with activities of 2 × 10^7^ Units/mg (PeproTech, Rehovot, Israel). Cells lines were then co-cultured with 100,000 (at density of 1×10^6^ cell/ml) IcAR-PD1 cells for 20–24 h. Supernatants were collected 20–24 h later, followed by testing with a murine IL-2 commercial sandwich ELISA kit (BioLegend).

### Generation of PDXs

Male NOD/SCID (Envigo-NOD.CB17-Prkdcscid/NCrHsd) mice were used for the study. Patient-derived tumor tissue samples were implanted subcutaneously into the dorsal flanks of the mice to form the PDXs. Tumor take rate varied from 1 to 6 months. PDXs were maintained by passaging the tumors in mice from first generation to subsequent generations. All animal experiments were conducted under the guidelines of the Institutional Animal Care and Use Committee (IACUC) of Ben-Gurion University of the Negev, according to specified protocols aiming to ensure animal welfare and reduce suffering (ethical clearance protocol number IL-80-12-2015).

### Fresh PDX co-culture

A PDX was removed when it reached approximately 500 mm^3^ in size. After aseptic excision of the PDXs, they were cut into 2 × 2 × 2 mm^3^ tissue explants. Each PDX sample was then co-cultured in a flat 96-well plate with 100,000 IcAR-PD1 cells (at density of 1×10^6^ cell/ml) for 24 h. The supernatants were collected 24 h later, followed by testing with a murine IL-2 commercial sandwich ELISA kit (BioLegend).

### FFPE co-culture

Five 5-micron FFPE sections were each placed in a 50-mL tube. The tissue sections were de-deparaffinized with HistoChoice^®^ (#H2779, Sigma-Aldrich) 3 times, 10 min each. Samples were then hydrated gradually an alcohol series: wash in 100% ethanol (twice, 5 min each), 95% ethanol (once, 5 min) and 70% ethanol (once, 5 min). Samples were then washed in deionized water (3 times, 5 min each). For antigen retrieval, samples were placed in 10 mL of Tris-EDTA 0.05% Tween buffer (10 mM, pH 9) and heated in a boiling water bath (95°C) for 30 min. Samples were then washed in deionized water (3 times, 5 min each). FFPE samples were then suspended in 1 mL of 1X PBS and seeded in a flat 96-well plate at 50 μL of sample per well. The culture plate was then dried in an oven (60 ºC for 1 h). Surface area coverage was determined with a JuliStage Recorder and analyzed with JuliStat software using the growth curve function to analyze the coverage (mm^2^/well) of samples in each well. Sample wells were then blocked overnight with 2% bovine serum albumin (BSA) in 1X PBS. For co-culture, 100,000 (at density of 1×10^6^ cell/ml) IcAR-PD1 cells were seeded on top of the FFPE samples for 24 h. Supernatants were collected 24 h later, followed by testing with a murine IL-2 commercial sandwich ELISA kit (BioLegend).

### Evaluation of IcAR activation via measurement of murine IL-2

Supernatants from co-culture of IcAR-PD1 and targets were collected after 20–24 h of incubation and analyzed for murine IL-2 by an ELISA assay. The 96-well ELISA plates were pre-coated with purified anti-mouse IL-2 (BioLegend) using coating buffer (0.1 M Na_2_HPO; pH 9.0), blocked with 10% FBS in PBST (0.05% Tween-20), and coated with the supernatant, followed by the addition of biotinylated anti-mouse IL-2 (BioLegend), and then streptavidin-horseradish peroxidase conjugate (SA-HRP) (Jackson ImmunoResearch, PA, USA) and 3,3’,5,5’-tetramethylbenzidine (TMB) (Dako, Denmark) for detection of murine IL-2.

## Statistical analysis

Statistical analyses were performed with GraphPad Prism8. Data in bar graphs are presented as means ± SEM. Correlation analysis was done using Pearson correlation assays, and significance was determined by two-way ANOVA or t-test (paired or unpaired). Differences were considered to be statistically significant at a two‐sided *P* < 0.05.

## Acknowledgments

We would like to thank the Oncology and Pathology Departments at Soroka University Medical Center (SUMC) and the Pathology Department at Barzilai Medical Center (BMC). We also want to thank the members of our laboratories for comments, suggestions and support during this work.

## Author contributions

Conceptualization: B.K., M.E. and A.P. Methodology: B.K. and O.A. Software: B.K. Resources: A.Y., D.A., M.S. and Y.G. Formal analysis: B.K. Visualization: B.K. Supervision: A.L., R.S.L, K.R., M.E. and A.P. Writing—original draft: B.K. Writing—review and editing: M.E. and A.P.

## Data and materials availability

All data are available in the main text or the supplementary materials.

## References

1. H. Borghaei, S. Gettinger, E. E. Vokes, L. Q. M. Chow, M. A. Burgio, J. de Castro Carpeno, A. Pluzanski, O. Arrietac, O. A. Frontera, R. Chiari, C. Butts, J. Wójcik-Tomaszewska, B. Coudert, M. C. Garassino, N. Ready, E. Felip, M. A. García, D. Waterhouse, M. Domine, F. Barlesi, S. Antonia, M. Wohlleber, D. E. Gerber, G. Czyzewicz, D. R. Spigel, L. Crino, W. E. E. Eberhardt, A. Li, S. Marimuthu, J. Brahmerc, Five-Year Outcomes From the Randomized, Phase III Trials CheckMate 017 and 057: Nivolumab Versus Docetaxel in Previously Treated Non–Small-Cell Lung Cancer. J. Clin. Oncol. 39, 723 (2021).

2. F. S. Hodi, V. Chiarion-Sileni, R. Gonzalez, J. J. Grob, P. Rutkowski, C. L. Cowey, C. D. Lao, D. Schadendorf, J. Wagstaff, R. Dummer, P. F. Ferrucci, M. Smylie, A. Hill, D. Hogg, I. Marquez-Rodas, J. Jiang, J. Rizzo, J. Larkin, J. D. Wolchok, Nivolumab plus ipilimumab or nivolumab alone versus ipilimumab alone in advanced melanoma (CheckMate 067): 4-year outcomes of a multicentre, randomised, phase 3 trial. Lancet Oncol. 19, 1480–1492 (2018).

3. T. S. K. Mok, Y. L. Wu, I. Kudaba, D. M. Kowalski, B. C. Cho, H. Z. Turna, G. Castro, V. Srimuninnimit, K. K. Laktionov, I. Bondarenko, K. Kubota, G. M. Lubiniecki, J. Zhang, D. Kush, G. Lopes, G. Gomez Aubin, L. Fein, D. Kaen, R. Kowalyszyn, G. Lerzo, G. Martinengo, M. Molina, E. Richardet, P. Picon, M. Varela, J. J. Zarba, S. J. de Azevedo, C. H. Barrios, C. Beato, C. A. S. Cerny, P. R. M. De Marchi, G. Fernandes, F. A. Franke, H. Freitas, G. Girotto, V. Lopes, L. Santos, M. A. Costa, A. K. Shimada, O. Smaletz, J. P. H. Soares, A. P. Victorino, C. Ferreira, M. Koleva, K. Koynov, R. Micheva, T. Deliverski, Z. Milanova, B. Doganov, S. Cheng, F. De Angelis, G. Speranza, R. A. Juergens, D. Ksienski, D. Fenton, O. Aren, C. Caglevic, H. Galindo, F. Rey, J. Chang, G. Chen, X. Chen, X. Ouyang, Y. Cheng, Z. Ding, M. Hou, Y. Fan, J. Feng, J. He, Y. He, Y. Hu, W. Li, X. Liu, Z. Liu, S. Lu, S. Qin, Q. Tang, B. Wang, K. Wang, L. Zhang, X. Zhang, J. Zhao, J. Wang, C. Zhou, J. Zhou, Q. Zhou, A. Cardona, R. Duarte, L. Gomez Wolff, A. Zambrano, M. Vallejo, L. Havel, V. Kolek, P. Kolman, L. Koubkova, L. Petruzelka, P. Popelkova, J. Roubec, J. Vanasek, T. Vlasek, J. Jaal, G. Kuusk, O. Avendano, H. Castro, K. Lopez, M. Sandoval, C. M. J. Ho, S. H. Lo, I. Laczo, B. Piko, G. Ostoros, K. Aoe, Y. Fujisaka, T. Hirashima, A. Horiike, Y. Hosomi, K. Hotta, M. Ichiki, F. Imamura, Y. Iwamoto, K. Kasahara, N. Katakami, T. Kato, S. Murakami, T. Kawaguchi, K. Kishi, T. Kurata, Y. Torii, Y. Nakahara, T. Nishimura, T. Ohira, H. Saka, T. Sawa, N. Seki, S. Sugawara, K. Takahashi, N. Takigawa, H. Tanaka, K. Yamada, T. Yokoyama, T. Yokoyama, H. Yoshioka, G. Purkalne, Z. Stara, A. Cesas, S. Cicenas, M. Zemaitis, S. H. How, C. K. Liam, C. K. Ong, L. M. Tho, O. Arrieta Rodriguez, F. de The Bustamante Valles, C. Hernandez Hernandez, L. Mas, L. Vera, J. Salas, H. Tejada, R. Edusma-Dy, C. Galvez, G. E. I. Ladrera, J. Tan Chun Bing, J. Jassem, E. Kalinka-Warzocha, B. Karaszewska, A. Kazarnowicz, K. Lesniewski Kmak, R. Ramlau, A. Araujo, F. Barata, N. Gil, V. Hespanhol, A. Alexandru, M. Dediu, N. Cherciu, D. Ciurescu, D. Ganea, L. Miron, D. Sirbu, M. Turdean, S. Emelyanov, N. Karaseva, L. Kuzina, S. Lazarev, I. Lifirenko, L. Bolotina, O. Lipatov, E. Ovchinnikova, M. Matrosova, A. Alyasova, A. Poltoratsky, P. Taranov, O. Zarubenkov, G. Cohen, L. Dreosti, F. Seolwane, J. Hall, G. Hart, C. Jordaan, S. Buddu, M. Botha, G. Landers, B. Rappaport, P. Ruff, L. Shepherd, W. Szpak, M. J. Ahn, J. H. Kim, P. Bergstrom, R. Ohman, H. Griph, D. Betticher, A. Ochsenbein, A. Zippelius, G. C. Chan, C. H. Chiu, T. C. Hsia, W. C. Su, C. H. Yang, T. Ativitavas, P. Danchaivijitr, K. Seetalarom, A. Sookprasert, V. Sriuranpong, O. Altundag, F. Cay Senler, M. Erman, T. Goksel, E. Goker, O. Ozyilkan, M. Seker, M. Gumus, F. Yumuk, G. Adamchuk, O. Ivashchuk, O. Ponomarova, A. Rusyn, S. Shevnya, Y. Shparyk, I. Sinielnikov, O. Andrusenko, D. Trukhyn, G. Ursol, I. Vynnychenko, T. Q. Nguyen, X. D. Pham, Pembrolizumab versus chemotherapy for previously untreated, PD-L1-expressing, locally advanced or metastatic non-small-cell lung cancer (KEYNOTE-042): a randomised, open-label, controlled, phase 3 trial. Lancet. 393, 1819–1830 (2019).

4. R. S. Herbst, P. Baas, D. W. Kim, E. Felip, J. L. Pérez-Gracia, J. Y. Han, J. Molina, J. H. Kim, C. D. Arvis, M. J. Ahn, M. Majem, M. J. Fidler, G. De Castro, M. Garrido, G. M. Lubiniecki, Y. Shentu, E. Im, M. Dolled-Filhart, E. B. Garon, Pembrolizumab versus docetaxel for previously treated, PD-L1-positive, advanced non-small-cell lung cancer (KEYNOTE-010): a randomised controlled trial. Lancet. 387, 1540–1550 (2016).

5. S. J. Antonia, A. Villegas, D. Daniel, D. Vicente, S. Murakami, R. Hui, T. Kurata, A. Chiappori, K. H. Lee, M. de Wit, B. C. Cho, M. Bourhaba, X. Quantin, T. Tokito, T. Mekhail, D. Planchard, Y.-C. Kim, C. S. Karapetis, S. Hiret, G. Ostoros, K. Kubota, J. E. Gray, L. Paz-Ares, J. de Castro Carpeño, C. Faivre-Finn, M. Reck, J. Vansteenkiste, D. R. Spigel, C. Wadsworth, G. Melillo, M. Taboada, P. A. Dennis, M. Özgüroğlu, Overall Survival with Durvalumab after Chemoradiotherapy in Stage III NSCLC. N. Engl. J. Med. 379, 2342–2350 (2018).

6. C. Faivre-Finn, D. Vicente, T. Kurata, D. Planchard, L. Paz-Ares, J. F. Vansteenkiste, D. R. Spigel, M. C. Garassino, M. Reck, S. Senan, J. Naidoo, A. Rimner, Y. L. Wu, J. E. Gray, M. Özgüroğlu, K. H. Lee, B. C. Cho, T. Kato, M. de Wit, M. Newton, L. Wang, P. Thiyagarajah, S. J. Antonia, Four-Year Survival With Durvalumab After Chemoradiotherapy in Stage III NSCLC—an Update From the PACIFIC Trial. J. Thorac. Oncol. 16, 860–867 (2021).

7. C. Robert, G. V Long, B. Brady, C. Dutriaux, M. Maio, L. Mortier, J. C. Hassel, P. Rutkowski, C. Mcneil, E. Kalinka-Warzocha, K. J. Savage, M. M. Hernberg, C. Lebbé, J. Charles, C. Mihalcioiu, V. Chiarion-Sileni, C. Mauch, F. Cognetti, A. Arance, H. Schmidt, D. Schadendorf, H. Gogas, L. Lundgren-Eriksson, C. Horak, B. Sharkey, I. M. Waxman, V. Atkinson, P. A. Ascierto, Comparative Analysis of Predictive Biomarkers for PD-1/PD-L1 Inhibitors in Cancers: Developments and Challenges. Cancers 2022, Vol. 14, Page 109. 14, 109 (2021).

8. S. Lu, J. E. Stein, D. L. Rimm, D. W. Wang, J. M. Bell, D. B. Johnson, J. A. Sosman, K. A. Schalper, R. A. Anders, H. Wang, C. Hoyt, D. M. Pardoll, L. Danilova, J. M. Taube, Comparison of Biomarker Modalities for Predicting Response to PD-1/PD-L1 Checkpoint Blockade: A Systematic Review and Meta-analysis. JAMA Oncol. 5, 1195–1204 (2019).

9. L. Marcus, S. J. Lemery, P. Keegan, R. Pazdur, FDA approval summary: Pembrolizumab for the treatment of microsatellite instability-high solid tumors. Clin. Cancer Res. 25, 3753–3758 (2019).

10. A. Marabelle, D. T. Le, P. A. Ascierto, A. M. Di Giacomo, A. de Jesus-Acosta, J. P. Delord, R. Geva, M. Gottfried, N. Penel, A. R. Hansen, S. A. Piha-Paul, T. Doi, B. Gao, H. C. Chung, J. Lopez-Martin, Y. J. Bang, R. S. Frommer, M. Shah, R. Ghori, A. K. Joe, S. K. Pruitt, L. A. Diaz, Efficacy of pembrolizumab in patients with noncolorectal high microsatellite instability/mismatch repair–deficient cancer: Results from the phase II KEYNOTE-158 study. J. Clin. Oncol. 38, 1–10 (2020).

11. A. A. Davis, V. G. Patel, The role of PD-L1 expression as a predictive biomarker: An analysis of all US food and drug administration (FDA) approvals of immune checkpoint inhibitors. J. Immunother. Cancer. 7, 1–8 (2019).

12. D. B. Doroshow, S. Bhalla, M. B. Beasley, L. M. Sholl, K. M. Kerr, S. Gnjatic, I. I. Wistuba, D. L. Rimm, M. S. Tsao, F. R. Hirsch, PD-L1 as a biomarker of response to immune-checkpoint inhibitors. Nat. Rev. Clin. Oncol. 2021 186. 18, 345–362 (2021).

13. A. A. Davis, V. G. Patel, The role of PD-L1 expression as a predictive biomarker: An analysis of all US food and drug administration (FDA) approvals of immune checkpoint inhibitors. J. Immunother. Cancer. 7, 1–8 (2019).

14. V. A. Boussiotis, P. Chatterjee, L. Li, Biochemical signaling of PD-1 on T cells and its functional implications. Cancer J. (United States). 20 (2014), pp. 265–271.

15. Y. Iwai, M. Ishida, Y. Tanaka, T. Okazaki, T. Honjo, N. Minato, Involvement of PD-L1 on tumor cells in the escape from host immune system and tumor immunotherapy by PD-L1 blockade. Proc. Natl. Acad. Sci. 99, 12293–12297 (2002).

16. Y. Latchman, C. R. Wood, T. Chernova, D. Chaudhary, M. Borde, I. Chernova, Y. Iwai, A. J. Long, J. A. Brown, R. Nunes, E. A. Greenfield, K. Bourque, V. A. Boussiotis, L. L. Carter, B. M. Carreno, N. Malenkovich, H. Nishimura, T. Okazaki, T. Honjo, A. H. Sharpe, G. J. Freeman, PD-L2 is a second ligand for PD-1 and inhibits T cell activation. Nat. Immunol. 2001 23. 2, 261–268 (2001).

17. J. Zhang, F. Dang, J. Ren, W. Wei, Biochemical Aspects of PD-L1 Regulation in Cancer Immunotherapy. Trends Biochem. Sci. 43, 1014–1032 (2018).

18. C. W. Li, S. O. Lim, W. Xia, H. H. Lee, L. C. Chan, C. W. Kuo, K. H. Khoo, S. S. Chang, J. H. Cha, T. Kim, J. L. Hsu, Y. Wu, J. M. Hsu, H. Yamaguchi, Q. Ding, Y. Wang, J. Yao, C. C. Lee, H. J. Wu, A. A. Sahin, J. P. Allison, D. Yu, G. N. Hortobagyi, M. C. Hung, Glycosylation and stabilization of programmed death ligand-1 suppresses T-cell activity. Nat. Commun. 7 (2016), doi:10.1038/ncomms12632.

19. K. J. Lastwika, W. Wilson, Q. K. Li, J. Norris, H. Xu, S. R. Ghazarian, H. Kitagawa, S. Kawabata, J. M. Taube, S. Yao, L. N. Liu, J. J. Gills, P. A. Dennis, Control of PD-L1 expression by oncogenic activation of the AKT-mTOR pathway in non-small cell lung cancer. Cancer Res. 76, 227–238 (2016).

20. J. H. Cha, L. C. Chan, C. W. Li, J. L. Hsu, M. C. Hung, Mechanisms Controlling PD-L1 Expression in Cancer. Mol. Cell. 76, 359–370 (2019).

21. C. M. Maher, J. D. Thomas, D. A. Haas, C. G. Longen, H. M. Oyer, J. Y. Tong, F. J. Kim, Small-Molecule Sigma1 Modulator Induces Autophagic Degradation of PD-L1. Mol. Cancer Res. 16, 243–255 (2018).

22. J. M. Pitt, M. Vétizou, R. Daillère, M. P. Roberti, T. Yamazaki, B. Routy, P. Lepage, I. G. Boneca, M. Chamaillard, G. Kroemer, L. Zitvogel, Resistance Mechanisms to Immune-Checkpoint Blockade in Cancer: Tumor-Intrinsic and -Extrinsic Factors. Immunity. 44, 1255–1269 (2016).

23. C. Sun, R. Mezzadra, T. N. Schumacher, Regulation and Function of the PD-L1 Checkpoint. Immunity. 48, 434–452 (2018).

24. D. M. Pardoll, The blockade of immune checkpoints in cancer immunotherapy. Nat. Rev. Cancer. 12, 252–264 (2012).

25. A. Haslam, J. Gill, V. Prasad, Estimation of the Percentage of US Patients With Cancer Who Are Eligible for Immune Checkpoint Inhibitor Drugs. JAMA Netw. open. 3, e200423 (2020).

26. C. Plazy, D. Hannani, E. Gobbini, Immune Checkpoint Inhibitor Rechallenge and Resumption: a Systematic Review. Curr. Oncol. Rep. (2022), doi:10.1007/S11912-022-01241-Z.

27. L. B. Kennedy, A. K. S. Salama, A review of cancer immunotherapy toxicity. CA. Cancer J. Clin. 70 (2020), pp. 86–104.

28. M. A. Postow, R. Sidlow, M. D. Hellmann, Immune-Related Adverse Events Associated with Immune Checkpoint Blockade. N. Engl. J. Med. 378 (2018), pp. 158–168.

29. M. K. Nesline, T. Knight, S. Colman, K. Patel, Economic Burden of Checkpoint Inhibitor Immunotherapy for the Treatment of Non–Small Cell Lung Cancer in US Clinical Practice. Clin. Ther. 42, 1682-1698.e7 (2020).

30. G. Morad, B. A. Helmink, P. Sharma, J. A. Wargo, Hallmarks of response, resistance, and toxicity to immune checkpoint blockade. Cell. 184, 5309–5337 (2021).

31. X. Wang, F. Teng, L. Kong, J. Yu, PD-L1 expression in human cancers and its association with clinical outcomes (2016), doi:10.2147/OTT.S105862.

32. Y. Lei, X. Li, Q. Huang, X. Zheng, M. Liu, Progress and Challenges of Predictive Biomarkers for Immune Checkpoint Blockade. Front. Oncol. 11, 609 (2021).

33. R. M. Samstein, C. H. Lee, A. N. Shoushtari, M. D. Hellmann, R. Shen, Y. Y. Janjigian, D. Barron, A. Zehir, E. J. Jordan, A. Omuro, T. J. Kaley, S. M. Kendall, R. J. Motzer, A. A. Hakimi, M. H. Voss, P. Russo, J. Rosenberg, G. Iyer, B. H. Bochner, D. F. Bajorin, H. A. Al-Ahmadie, J. E. Chaft, C. M. Rudin, G. J. Riely, S. Baxi, A. L. Ho, R. J. Wong, D. G. Pfister, J. D. Wolchok, C. A. Barker, P. H. Gutin, C. W. Brennan, V. Tabar, I. K. Mellinghoff, L. M. DeAngelis, C. E. Ariyan, N. Lee, W. D. Tap, M. M. Gounder, S. P. D’Angelo, L. Saltz, Z. K. Stadler, H. I. Scher, J. Baselga, P. Razavi, C. A. Klebanoff, R. Yaeger, N. H. Segal, G. Y. Ku, R. P. DeMatteo, M. Ladanyi, N. A. Rizvi, M. F. Berger, N. Riaz, D. B. Solit, T. A. Chan, L. G. T. Morris, Tumor mutational load predicts survival after immunotherapy across multiple cancer types. Nat. Genet. 51 (2019), pp. 202–206.

34. R. Cristescu, D. Aurora-Garg, A. Albright, L. Xu, X. Q. Liu, A. Loboda, L. Lang, F. Jin, E. H. Rubin, A. Snyder, J. Lunceford, Tumor mutational burden predicts the efficacy of pembrolizumab monotherapy: a pan-tumor retrospective analysis of participants with advanced solid tumors. J. Immunother. cancer. 10 (2022), doi:10.1136/JITC-2021-003091.

35. R. Cristescu, R. Mogg, M. Ayers, A. Albright, E. Murphy, J. Yearley, X. Sher, X. Q. Liu, H. Lu, M. Nebozhyn, C. Zhang, J. K. Lunceford, A. Joe, J. Cheng, A. L. Webber, N. Ibrahim, E. R. Plimack, P. A. Ott, T. Y. Seiwert, A. Ribas, T. K. McClanahan, J. E. Tomassini, A. Loboda, D. Kaufman, Pan-tumor genomic biomarkers for PD-1 checkpoint blockade-based immunotherapy. Science (80-.). 362 (2018), doi:10.1126/science.aar3593.

36. J. S. Lee, E. Ruppin, Multiomics Prediction of Response Rates to Therapies to Inhibit Programmed Cell Death 1 and Programmed Cell Death 1 Ligand 1. JAMA Oncol. 5, 1614–1618 (2019).

37. K. Wang, S. Patkar, J. S. Lee, E. M. Gertz, W. Robinson, F. Schischlik, D. R. Crawford, A. Schäffer, E. Ruppin, Deconvolving Clinically Relevant Cellular Immune Cross-talk from Bulk Gene Expression Using CODEFACS and LIRICS Stratifies Patients with Melanoma to Anti–PD-1 Therapy. Cancer Discov. 12, 1088–1105 (2022).

38. V. Lazar, S. Magidi, N. Girard, A. Savignoni, J. F. Martini, G. Massimini, C. Bresson, R. Berger, A. Onn, J. Raynaud, F. Wunder, I. Berindan-Neagoe, M. Sekacheva, I. Braña, J. Tabernero, E. Felip, A. Porgador, C. Kleinman, G. Batist, B. Solomon, A. M. Tsimberidou, J. C. Soria, E. Rubin, R. Kurzrock, R. L. Schilsky, Digital Display Precision Predictor: the prototype of a global biomarker model to guide treatments with targeted therapy and predict progression-free survival. NPJ Precis. Oncol. 5 (2021), doi:10.1038/S41698-021-00171-6.

39. E. Sterner, N. Flanagan, J. C. Gildersleeve, Perspectives on Anti-Glycan Antibodies Gleaned from Development of a Community Resource Database. ACS Chem. Biol. 11, 1773–1783 (2016).

40. X. Bu, V. R. Juneja, C. G. Reynolds, K. M. Mahoney, M. T. Bu, K. A. McGuire, S. Maleri, P. Hua, B. Zhu, S. R. Klein, E. A. Greenfield, P. Armand, J. Ritz, A. H. Sharpe, G. J. Freeman, Monitoring PD-1 Phosphorylation to Evaluate PD-1 Signaling during Antitumor Immune Responses. Cancer Immunol. Res. 9, 1465–1475 (2021).

41. Y. N. Wang, H. H. Lee, J. L. Hsu, D. Yu, M. C. Hung, The impact of PD-L1 N-linked glycosylation on cancer therapy and clinical diagnosis. J. Biomed. Sci. 27, 1–11 (2020).

42. H. H. Lee, Y. N. Wang, W. Xia, C. H. Chen, K. M. Rau, L. Ye, Y. Wei, C. K. Chou, S. C. Wang, M. Yan, C. Y. Tu, T. C. Hsia, S. F. Chiang, K. S. C. Chao, I. I. Wistuba, J. L. Hsu, G. N. Hortobagyi, M. C. Hung, Removal of N-Linked Glycosylation Enhances PD-L1 Detection and Predicts Anti-PD-1/PD-L1 Therapeutic Efficacy. Cancer Cell. 36, 168-178.e4 (2019).

43. M. Wang, S. Wang, J. A. Trapani, P. J. Neeson, Challenges of PD-L1 testing in non-small cell lung cancer and beyond. J. Thorac. Dis. 12, 4541 (2020).

44. D. Agilent Technologies, PD-L1 IHC 22C3 pharmDx Interpretation Manual – NSCLC. Dako, 1–76 (2018).

45. H. Brunnström, A. Johansson, S. Westbom-Fremer, M. Backman, D. Djureinovic, A. Patthey, M. Isaksson-Mettävainio, M. Gulyas, P. Micke, PD-L1 immunohistochemistry in clinical diagnostics of lung cancer: Inter-pathologist variability is higher than assay variability. Mod. Pathol. 30, 1411–1421 (2017).

46. Y. R. Miao, K. N. Thakkar, J. Qian, M. S. Kariolis, W. Huang, S. Nandagopal, T. T. Chi Yang, A. N. Diep, G. M. Cherf, Y. Xu, E. J. Moon, Y. Xiao, H. Alemany, T. Li, W. Yu, B. Wei, E. B. Rankin, A. J. Giaccia, Neutralization of PD-L2 is Essential for Overcoming Immune Checkpoint Blockade Resistance in Ovarian Cancer. Clin. Cancer Res. 27, 4435–4448 (2021).

47. T. Tanegashima, Y. Togashi, K. Azuma, A. Kawahara, K. Ideguchi, D. Sugiyama, F. Kinoshita, J. Akiba, E. Kashiwagi, A. Takeuchi, T. Irie, K. Tatsugami, T. Hoshino, M. Eto, H. Nishikawa, Immune Suppression by PD-L2 against Spontaneous and Treatment-Related Antitumor Immunity. Clin. Cancer Res. 25, 4808–4819 (2019).

48. Y. Latchman, C. R. Wood, T. Chernova, D. Chaudhary, M. Borde, I. Chernova, Y. Iwai, A. J. Long, J. A. Brown, R. Nunes, E. A. Greenfield, K. Bourque, V. A. Boussiotis, L. L. Carter, B. M. Carreno, N. Malenkovich, H. Nishimura, T. Okazaki, T. Honjo, A. H. Sharpe, G. J. Freeman, PD-L2 is a second ligand for PD-1 and inhibits T cell activation. Nat. Immunol. 2, 261–268 (2001).

49. C. B. M. Du Puch, M. Vanderstraete, S. Giraud, C. Lautrette, N. Christou, M. Mathonnet, Benefits of functional assays in personalized cancer medicine: more than just a proof-of-concept. Theranostics. 11, 9538 (2021).

50. J. A. Marin-Acevedo, E. M. O. Kimbrough, Y. Lou, Next generation of immune checkpoint inhibitors and beyond. J. Hematol. Oncol. 14 (2021), doi:10.1186/S13045-021-01056-8.

51. L. Kraehenbuehl, C.-H. Weng, S. Eghbali, J. D. Wolchok, T. Merghoub, Enhancing immunotherapy in cancer by targeting emerging immunomodulatory pathways. Nat. Rev. Clin. Oncol. 2021, 1–14 (2021).

52. E. Ruffo, R. C. Wu, T. C. Bruno, C. J. Workman, D. A. A. Vignali, Lymphocyte-activation gene 3 (LAG3): The next immune checkpoint receptor. Semin. Immunol. 42 (2019), p. 101305.

53. P. S. Hegde, D. S. Chen, Top 10 Challenges in Cancer Immunotherapy. Immunity. 52 (2020), pp. 17–35.

54. D. A. Fabrizio, T. J. George, R. F. Dunne, G. Frampton, J. Sun, K. Gowen, M. Kennedy, J. Greenbowe, A. B. Schrock, A. F. Hezel, J. S. Ross, P. J. Stephens, S. M. Ali, V. A. Miller, M. Fakih, S. J. Klempner, Beyond microsatellite testing: assessment of tumor mutational burden identifies subsets of colorectal cancer who may respond to immune checkpoint inhibition. J. Gastrointest. Oncol. 9, 610 (2018).

