## Supplemental material for "Functional binding of PD1 ligands predicts response to anti-PD1 treatment in cancer patients"

Bar Kaufman *et al.*

**This PDF file includes:**

Figs. S1 to S4

Tables S1 to S2

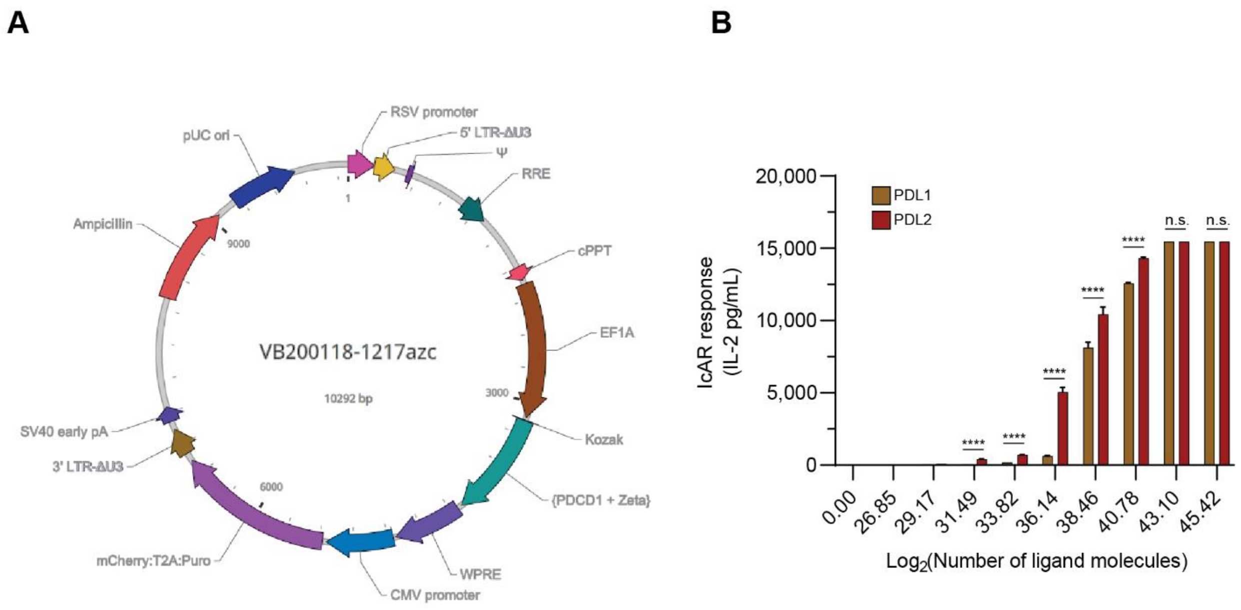


**Fig. S1. Generation of IcAR-PD1 cell line**

(A) IcAR-PD1 plasmid, consists of ectodomain of PDCD1 gene fused to transmembrane and intracellular domain of murine CD3ζ chain. Selection markers of infection consist of mCherry fluorophore and puromycin antibody resistance. (C) Response of IcAR-PD1 to ascending concentrations of rhPDL1 and rhPDL2, pre-coated in a 96-well plate. Concentration of coated ligands are shown as Log2 of the total number of ligand molecules coated in each well. (3 repetitions). ****$P$<0.0001.

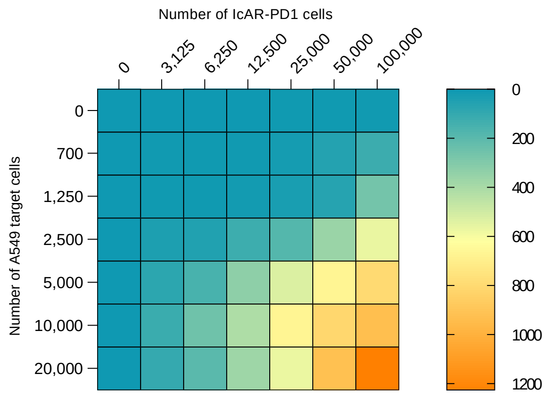


**Fig. S2. Determination of the number of effector cells**

A double titration of cytokine-induced target cells (A549) and IcAR-PD1 cells was performed. The titration ranged from 0 cells to 100,000 reporter cells and 20,000 target cells. At 100,000 reporter cells, IcAR-PD1 was able to respond to 700 target cells while still discriminating between different amounts of target cells. Results are shown as an average of three biological repeats.

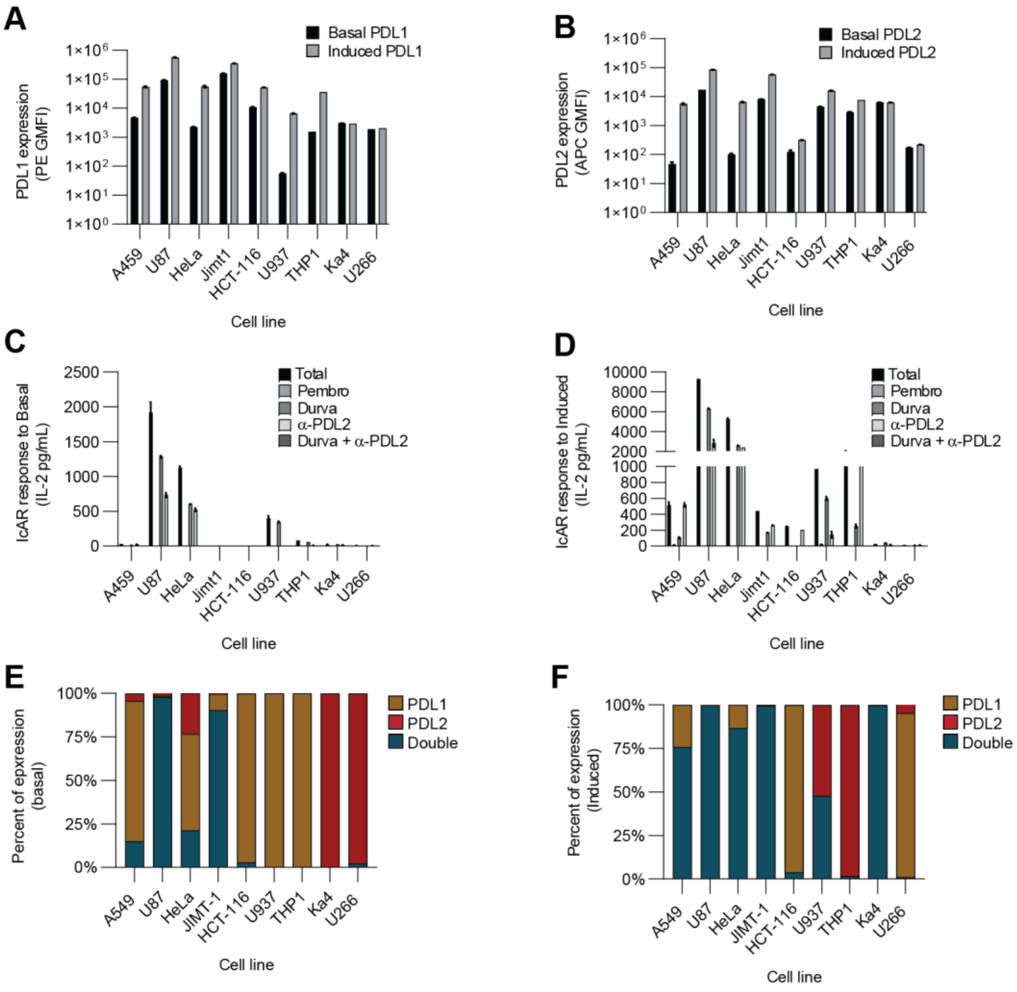


**Fig. S3. Ligand expression and IcAR response to cancer-cell lines**

(A) PDL1 expression of 9 different cell lines in their basal state (black) and their cytokine-induced state (gray). (B) PDL2 expression of different cell lines in their basal state (black) and their cytokine-induced state (gray). (C) IcAR-PD1 response (IL-2) to cell lines in their basal state: total response (black) and response after exposure to antibodies (Pembro = anti-PD1, Durva = anti-PDL1; different shades of gray). (D) IcAR-PD1 response (IL-2) to cell lines post induction with cytokines: total response (black) and response after exposure to antibodies (Pembro = anti-PD1, Durva = anti-PDL1; different shades of gray). (E) Percent of ligand expression in cell lines in basal state. Relative expression of each ligand was determined by flow-cytometry staining for both ligands. Events for each ligand were divided by the total number of positive cells (for both ligands) to calculate percent expression for each ligand.
(F) Percent of ligand expression in cell lines in their cytokine-induced state.

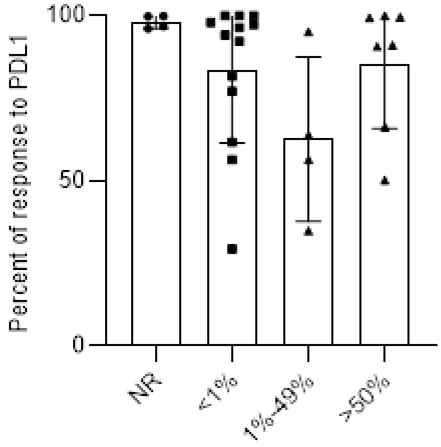


**Fig. S4. Percent of IcAR response to PDL1 as function of pathological grading of PDL1 (n=29).**

**Table S1.**

Cell line data

| Cell line | Origin tissue | Cell type | Disease | Properties |
|---|---|---|---|---|
| A549 | Lung | Epithelial cells | Carcinoma | Adherent |
| U87MG | Brain | Epithelial cells | Glioblastoma | Adherent |
| U937 | Pleural effusion | Monocyte | Histiocytic lymphoma | Suspension |
| HCT-116 | Large intestine; Colon | Epithelial cells | Colorectal carcinoma | Adherent |
| HeLa | Uterus; Cervix | Epithelial cells | Adenocarcinoma | Adherent |
| JIMT1 | Breast | Epithelial cells | Invasive ductal carcinoma | Adherent |
| 721.221 | Peripheral blood | B lymphocyte | Transformed cells line | Suspension |
| U266 | Peripheral blood | B lymphocyte | Myeloma | Suspension |
| THP-1 | Peripheral blood | Monocyte | Acute monocytic leukemia | Suspension |

**Table S2.**

PDX data

| PDX number | Pathology | PDX | Date of surgery/ implantation | Pathological evaluation of PDL1 staining | Treatment |
|---|---|---|---|---|---|
| 1 | Lung | LSE19 | 22.01.19 | %1> | Atezolizumab + Chemo/EGFR |
| 2 | Lung | LSE16 | 14.10.18 | %50< | Keytruda + chemo |
| 3 | Lung | LTE37 | 10.10.19 | %49-1 | Patient died one week post surgery |
| 4 | Lung | LSE20 | 31.03.19 | %1> | Keytruda+ chemo / nivolumab+ ipilimumab |
| 5 | HNSCC | HN6022 | 17.01.20 | %1> | Unknown |
